# Accurate prediction of orthologs in the presence of divergence after duplication

**DOI:** 10.1101/294405

**Authors:** Manuel Lafond, Mona Meghdari Miardan, David Sankoff

## Abstract

**Motivation:** When gene duplication occurs, one of the copies may become free of selective pressure and evolve at an accelerated pace. This has important consequences on the prediction of orthology relationships, since two orthologous genes separated by divergence after duplication may differ in both sequence and function. In this work, we make the distinction between the *primary* orthologs, which have not been affected by accelerated mutation rates on their evolutionary path, and the *secondary* orthologs, which have. Similarity-based prediction methods will tend to miss secondary orthologs, whereas phylogeny-based methods cannot separate primary and secondary orthologs. However, both types of orthology have applications in important areas such as gene function prediction and phylogenetic reconstruction, motivating the need for methods that can distinguish the two types.

**Results:** We formalize the notion of divergence after duplication, and provide a theoretical basis for the inference of primary and secondary orthologs. We then put these ideas to practice with the HyPPO (Hybrid Prediction of Paralogs and Orthologs) framework, which combines ideas from both similarity and phylogeny approaches. We apply our method to simulated and empirical datasets, and show that we achieve superior accuracy in predicting primary orthologs, secondary orthologs and paralogs.

**Availability:** HyPPO is a modular framework with a core developed in Python, and is provided with a variety of C_++_ modules. The source code is available at https://github.com/manuellafond/HyPPO.

## 1 Introduction

During the course of evolution, speciation and duplication create pairs of homologous genes, which can be classified into two categories. Two genes are orthologs if they descend from an ancestral gene that has undergone speciation, and paralogs if they result from duplication. Distinguishing orthologs from paralogs is of considerable importance in biology, owing to their functional and evolutionary implications (Koonin, 2005; Gabaldón and Koonin, 2013). Notably, the well-known “orthologs conjecture” states that orthologs tend to perform similar functions, in contrast with paralogs, which tend to diverge in their functional role. The reasoning underlying this conjecture is that when a gene creates a copy in the same genome, then this genome has two genes that are able to perform the same function. Hence, one of these copies may become free of selective pressure, resulting in a higher rate of mutations and facilitating a possible change in functionality. We refer the reader to (Studer and Robinson-Rechavi, 2009; Nehrt *et al.*, 2011; Altenhoff *et al.*, 2012; Thomas *et al.*, 2012; Chen and Zhang, 2012) for some recent developments on this conjecture.

The two traditional methods to predict orthology relations are *similarity-based* and *phylogeny-based* (see (Kristensen *et al.*, 2011; Altenhoff and Dessimoz, 2012) for a survey). Similarity-based methods aim to partition the set of genes into “groups” of orthologs, under the assumption that genes that share similar sequences are orthologous. The precise notion of a group depends on the particular method used, since in some cases, these groups include multiple genes from the same species (e.g. recent paralogs). For instance in OrthoMCL (Li *et al.*, 2003), Markov clustering is used to identify groups of orthologs and close paralogs. A similar idea is implemented in Proteinortho using spectral clustering techniques (Lechner *et al.*, 2011). The OMA software is somewhat more stringent, and aims to find cliques of pairwise orthologous genes (Roth *et al.*, 2008). This was extended in later versions with the addition of the GETHOGS algorithm (Altenhoff *et al.*, 2013; Train *et al.*, 2017). The program is able to potentially find all orthologous gene pairs by grouping genes into *hierarchical orthologous groups* (HOGs), which are the orthologs that belong to a set of species within a given taxonomic range. Some other similarity-based grouping methods and databases include COG (Tatusov *et al.*, 2003), EggNOG (Powell *et al.*, 2011), InParanoid (O’brien *et al.*, 2005) and OrthoFinder (Emms and Kelly, 2015). The OMG approach (Zheng *et al.*, 2011) identifies sets of pairwise *orthogonal* genes from a homology graph inferred from synteny.

On the other hand, phylogeny-based methods do not attempt in the first instance to group genes together, but rather aim to identify orthologs by inferring the lowest common ancestor (lca) relationships between genes directly. This can be done by inferring a gene tree and identifying its speciation and duplication events using *reconciliation* with a species tree (Stolzer *et al.*, 2012; Ullah *et al.*, 2015). This approach is however sensitive to errors in the gene tree and the species tree. Methods of this type include notably LOFT (Van der Heijden *et al.*, 2007) and COCO-CL (Jothi *et al.*, 2006). Recently, a number of methods exploit a fundamental property of orthology graphs, in which vertices are genes and edges depict orthology: a graph G is a valid orthology graph if and only if it is *P*__4__-free (Hernandez-Rosales *et al.*, 2012; Hellmuth *et al.*, 2013; Lafond and El-Mabrouk, 2014; Hellmuth *et al.*, 2015; Dondi *et al.*, 2017a; Lafond *et al.*, 2016; Jones *et al.*, 2016; Dondi *et al.*, 2017b), i.e. *G* has no path on 4 vertices without a shortcut. The essential reason underlying this characterization is that *P*_4_-free graphs have a special tree representation called a *co-tree*, which turns out to be interpretable as a gene tree that depicts all the lca relationships prescribed by *G*. These methods usually construct an approximate putative orthology graph and perform a minimum number of modifications on the graph so that it becomes *P*_4_-free.

The output from similarity-based and phylogeny-based methods can be different, as some orthologies may be irrelevant from the perspective of grouping genes with similar functions, but will be inferred by phylogenetic methods. Consider the example of Figure 1. The node *x* represents an ancestral duplication event in which *neofunctionalization* hypothetically occurred, where one copy (here the left child of *x*) retains its original functionality whereas the other copy (the right child of *x*) diverges and starts accumulating mutations at ahigher rate. Asa result of this event, the *a* and *b* genes remain similar, as no major divergence event separates the two. However, *a* and *c* will have differentiated significantly, even though the two genes are orthologous (since their lca is a speciation). Consequently, similarity-based methods are unlikely to mark *a* and *c* as orthologs, as the two genes do not share sequence similarity. From the point of view of functional annotation, this is not problematic, since the *a*, *c* orthology pair essentially behave as paralogs and may differ in function.

**Fig. 1:**
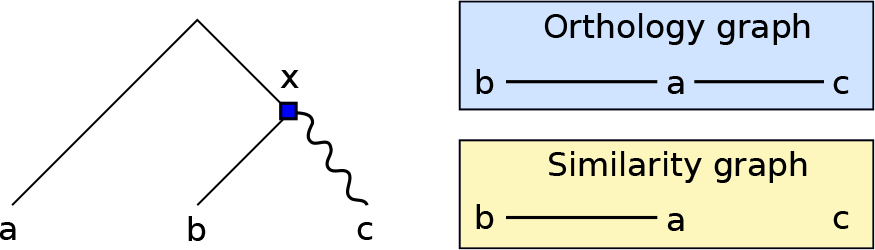
An example of gene tree for the gene family {*a*, *b*, *c*}, along with the underlying orthology and similarity graphs. The root of the tree is a speciation, the square is a duplication, and the wiggly edge represents an event of divergence after duplication. The gene pairs *ab* and *ac* are orthologs. However, *a* and *c* will not appear as “similar”, as there was a significant divergence event on their evolutionary path.

This raises the question as to which type of inference is the most relevant: should we focus on the groups of similar orthologs, or do we need *all* the orthologs? Ideally, both types of orthologies should be predicted, since similarity groups are useful for function-related applications (Doyle *et al.*, 2010), while the complete set of pairwise relations is useful for phylogenetic reconstructions (Hellmuth *et al.*, 2015).

Of course, post-duplication phenomena other than neofunctionalization as described above can arise. For instance, *bifunctionalization* occurs when both copies preserve the parental function without diverging, and *subfunctionalization* occurs when the functions of the common ancestor are partitioned among the descending paralogs (Zhang, 2003), in which case it is possible for both copies to undergo a new rate of mutations. Nevertheless, the phenomenon of a single copy undergoing functional change has been reported to occur frequently (Lynch and Conery, 2000; Jordan *et al.*, 2004; Woods *et al.*, 2013; Soria *et al.*, 2014; Cardoso-Moreira *et al.*, 2016), as duplication models other than neofunctionalization, for instance *adaptive radiation* or *modified duplication*, also predict this type of divergence after duplication. We redirect the reader to (Innan and Kondrashov, 2010, Table 1) for more details.

**Table 1.** Average accuracy, precision, recall and cluster scores, taken over the set of all simulated gene trees, for every possible value of *ℓ*.

|  | Accuracy | Precision | Recall | Cluster score |
| --- | --- | --- | --- | --- |
| HyPPO | .940 | .911 | .875 | .915 |
| HyPPO + Species Tree | .949 | .924 | .905 | .915 |
| OMA-GETHOGS | .877 | .940 | .699 | .831 |
| OrthoMCL | .812 | .845 | .496 | .690 |

In this work, we are interested in how the task of predicting orthology is affected when divergence after duplication does occur. We propose an orthology prediction framework that makes the distinction between the *primary orthologs*, the orthologous gene pairs that have not been separated by an event of duplication followed by an increased rate of mutation, and the *secondary orthologs*, which consist of the pairs of orthologs which have had at least one such event along their evolutionary path. For example in Figure 1, {*a*, *b*} are primary orthologs whereas {*a*, *c*} are secondary orthologs. Note that the relevance of making this distinction was also discussed in (Swenson and El-Mabrouk, 2012), where the primary orthologs are called *isoorthologs*. Also observe that this categorization is different from the co-orthology relationship, which is usually defined with respect to a given node of a gene tree. We formalize this notion of primary orthology and show that they must form cliques in the orthology graph, thereby providing a formal justification of the clustering steps often performed by orthology prediction methods.

Our algorithmic framework is a hybrid approach that makes use of both similarity clustering and phylogenetic reconstruction. More specifically, we use exact graph partitioning techniques to find the primary orthologs, construct a species tree from the orthology clusters found, and use this species tree to obtain the secondary orthologs. The method also puts to practical use the theory of *P*_4_-free orthology graphs developed recently. The framework, which we call HyPPO (for Hybrid Prediction of Paralogs and Orthologs), is implemented as a fully modular pipeline in which each task can have its own independent implementation.

We evaluate HyPPO on both simulated and real empirical datasets. We compare our method with OrthoMCL and OMA-GETHOGS - to the best of our knowledge, the latter is the only other program that can distinguish primary and secondary orthologs. We show that HyPPO achieves superior accuracy in the set of predicted pairwise relations, and is better at finding the primary orthology clusters.

## 2 Methods

In this section, we first introduce the preliminary notions necessary for the understanding of this paper. We then formalize the notion of divergence after duplication in gene trees and its consequences on distinguishing primary and secondary orthologs. We then present the main algorithmic components of the HyPPO framework. Note that due to space constraints, most of the proofs are relegated to the supplementary material.

### 2.1 Preliminary notions

In this paper, every tree *T* is assumed to be rooted and binary, i.e. each internal node has two children, and each node *v* has a unique parent *p*(*v*), with the exception of the root, denoted *r*(*T*), which has no parent. The set of vertices of *T* is denoted by *V*(*T*) and its set of leaves by *L*(*T*). A node *u* of *T* is an *ancestor* of a node *v* if *u* is on the path from *v* to *r*(*T*). Then *v* is a *descendant* of *u*, in which case we write *v* ≤ *u*. If *v* ≠ *u*, we write *v* < *u*. If none of *v* ≤ *u* nor *u* ≤ *v* holds, then *u* and *v* are *incomparable*, and we write *u* ⊕ *v*. The *least common ancestor* of a set of leaves *X* ∈ *L*(*T*), denoted *lca*_*T*_(*X*), is the node *u* of *T* that is the ancestor of every node in *X*, and that is the most distant from the root. For convenience, we may write *lca*_*T*_ (*a*, *b*) instead of *lca*_*T*_ ({*a*, *b*}). For a graph *G*, a *clique* is a set of vertices of *G* in which each pair shares an edge. A *P*_*k*_ is a path on *k* vertices with no chord.

A *gene family* Γ is a set of genes related by homology. A *gene tree* for Γ is a tree *T* in which *L*(*T*) = Γ. Likewise, if ∑ is a set of species, a *species tree* for ∑ is a tree *S* satisfying *L*(*S*) = ∑. Each gene *g* ∈ Γ belongs to a species denoted *σ*(*g*). If *X* ⊆ Γ is a set of genes, then we write *σ*(*X*) = {*σ*(*g*): *g* ∈ *X*}.

A DS-tree (*T*, *ℓ*) is a pair in which *T* is a gene tree, and *ℓ*: *V*(*T*) \ *L*(*T*) → {𝔻, 𝕊} is a function labeling internal nodes by either 𝔻 or 𝕊, respectively standing for duplication and speciation. We will often omit mentioning *ℓ* explicitly, and say that *T* is a *DS*-tree with the understanding that its internal nodes are labeled by *ℓ*. Given a species tree *S*, we map each node of *T* to a node of *S* with the *lca-mapping s*: *V*(*T*) → *V*(*S*) defined as follows: if *g* ∈ *L*(*T*), then *s*(*g*) = *σ*(*g*), and otherwise *s*(*g*) = *lca*_*s*_({*σ*(*l*): *l* ∈ *L*(*T*) and *l* ≤ *g*}). By a slight abuse of notation, if *C* ⊆ Γ, we denote *s*(*C*) = *lca*_*S*_(*σ*(*C*)). Note that *s* depends on the species tree *S*. In case of ambiguity, we may write *s*(*g*, *S*) (or *s*(*C*, *S*)) to denote the lca-mapping with respect to *S*. An example of lca-mapping is presented later on in Figure 3.

**Fig. 2.**
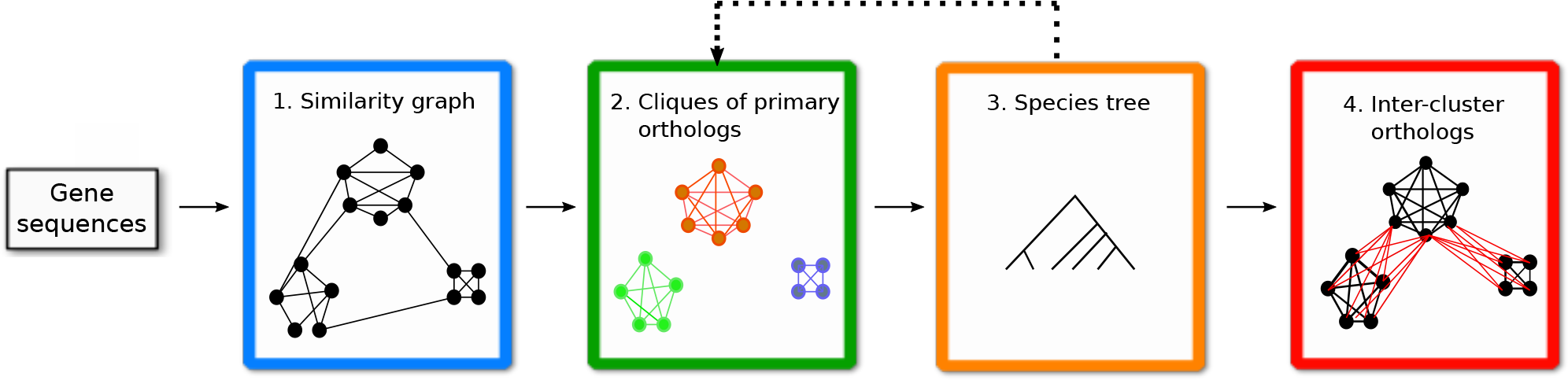
The four essential steps of the HyPPO pipeline. First, a similarity graph is constructed from the gene sequences. This graph is refined into orthology clusters, which are cliques of primary orthologs. These clusters are used to infer a species tree, which is then used to find the secondary orthologs (which we also call inter-cluster orthologs).

**Fig. 3:**
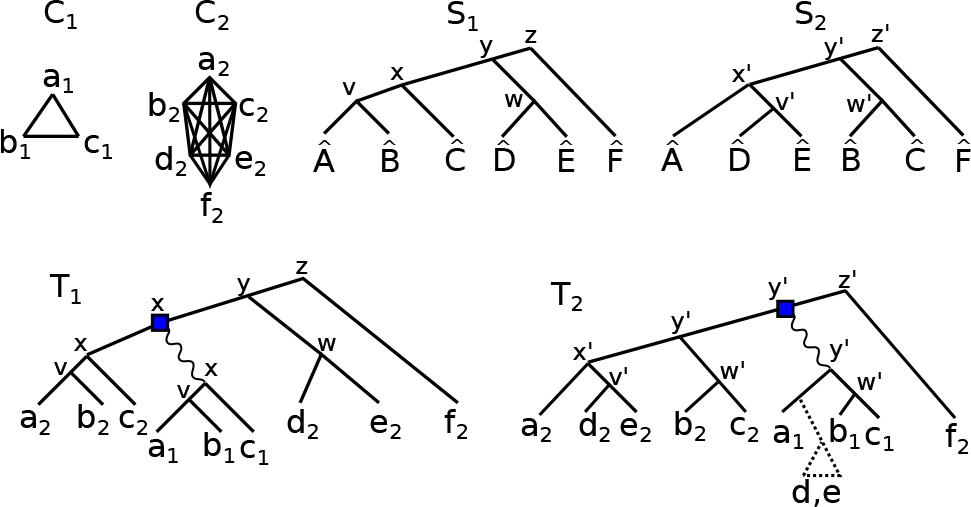
Two orthology clusters *C*_1_ and *C*_2_, and two putative species trees *S*_1_ and *S*_2_. The genes are named according to their species, e.g. *σ*(*a*_1_) = *Â*. Each internal node *u* of *T*_1_ (resp. *T*_2_) is labeled by its lca-mapping *s*(*u*, *S*_1_) (resp. *s*(*u*, *S*_2_)). The tree *T*_1_ is the DAD-tree that could have given rise to *C*_1_ and *C*_2_ if *S*_1_ was the true species tree, assuming that *C*_1_ was born during the history of the *C*_2_ genes. *T*_2_ is the DAD-tree for *C*_1_ and *C*_2_ under the assumption that *S*_2_ is the true species tree. In the case of *T*_2_, a loss in the clade 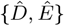 is required to explain the birth of the *C*_1_ cluster at a time prior to *lca*_S2_ (*σ*(*C*_1_)) = *y*′.

We use the definition of Fitch (Fitch, 2000) for orthology and paralogy.

#### Definition 1

*With respect to a DS-tree T, two genes a and b are* orthologs *if ℓ*(*lca*_*T*_(*a*, *b*)) = 𝕊, *and* paralogs *if ℓ*(*lca*_*T*_(*a*, *b*)) = 𝔻.

An *orthology graph G* = (*V*, *E*) is a graph in which *V* is a set of genes and *uv* ∈ *E* if and only if *u* and *v* are orthologs. An *orthology cluster C* is a set of genes that are pairwise orthologous. In particular, *C* may contain at most one gene from any given species.

### 2.2 Orthology and divergence after duplication

As mentioned in the introduction, duplication may introduce divergence when one of the child copies is redundant. In this section, our goal is to investigate the implications of this phenomenon on the structure of orthology relations. That is, if divergence after duplication occurs, what are the orthologs that are expected to be recovered using similarity-based methods? Or using phylogeny-based methods? We present the tools that we will use to devise methods tailored for this type of evolution. As we will demonstrate in the Results section, the methodology developed here can also be used in case that divergence after duplication does not always occur, or even when if it does not happen at all.

We formalize divergence after duplication as follows. Let *T* be a *DS*-tree. Suppose that for each 𝔻 node *v* of *T*, *v* has one child *v*_1_ that evolved at a normal rate, and another child *v*_2_ that underwent an accelerated pace of evolutionary changes. We will say that the *vv*_2_ edge is a *divergent duplication edge*. Now, we will assume that if, for two leaves *g*_1_ and *g*_2_ of *L*(*T*), there is a divergent duplication edge on the path between *g*_1_ and *g*_2_, then *g*_1_ and *g*_2_ will be considered “not similar”. Conversely, we will assume that if there is no such edge on the path between *g*_1_ and *g*_2_, then no unexpected differentiation between *g*_1_ and *g*_2_ will be observed, and thus *g*_1_ and *g*_2_ will be considered “similar”.

Note that here, we have not defined “similarity” in a quantitative manner. For our purposes, we prefer to leave this notion abstract, and we will assume that divergence after duplication leaves behind some traces that can be used to detect similarity versus non-similarity (for instance, one could compare the sequence of a gene to all its homologs in a given species). This is because our goal is to evaluate the consequences of divergence in the orthology inference process, and not necessarily to detect such divergence. We thus prefer to keep the notion of similarity as general as possible. We do note however that in the future, further investigation will be needed to establish concrete methods to recognize similarity.

We now introduce DAD-trees (for Divergence After Duplication).

#### Definition 2

*A DAD-tree T for* Γ *is a DS-tree for* Γ *in which each* 𝔻 *node v has exactly one child v′ such that the edge vv′ is marked as divergent, and the other edges of T are marked as non-divergent*.

*Two genes g*_1_, *g*_2_ ∈ *L*(*T*) *are called* similar *if there is no divergent edge on the path between g*_1_ *and g*_2_ *in T*. *Otherwise, g*_1_ *and g*_2_ *are called* non-similar.

It then becomes quite easy to characterize similarity graphs, using the above notion of “similarity”.

#### Proposition 3

*Let G be the graph in which V*(*G*) = Γ, *and two genes share an edge if and only if they are similar with respect to a DAD-tree T. Then each connected component of G is a clique*.

Therefore, unlike orthology, this notion of similarity is a transitive relation. Proposition 3 then tells us precisely what we should expect from similarity graphs when divergence after duplication always occurs: we should obtain cliques of genes. Moreover, the genes in these cliques are pairwise orthologous, as two paralogous genes cannot appear as “similar” under the above definition. Hence, the primary orthologs resulting from a DAD-tree form the cliques in the similarity graph. For this reason, we may sometimes call *orthology clusters* the cliques of primary orthologs. This motivates the first essential step of our orthology prediction framework, which is to transform the similarity graph into a cluster graph. However, this will only identify a subset of all orthologies, as there may be inter-cluster orthologies.

One idea might be to infer orthology between the “closest” inter-cluster genes. However, this may introduce invalid relations or inconsistencies. Here, as in (Hellmuth *et al.*, 2015; Lafond *et al.*, 2016), we take the viewpoint that for a set of relations to be correct, there must be an evolutionary scenario that could have given rise to the observed relations. In other words, there should exist a *DS*-tree that displays the inferred orthologies, and the speciation nodes on this *DS*-tree should agree with the species history. This notion of agreement with a species tree is defined as in (Lafond and El-Mabrouk, 2014; Jones *et al.*, 2016).

#### Definition 4

*Let T be a DS-tree and S be a species tree. Moreover, let v* ∈ *V*(*T*) \ *L*(*T*) *be an* 𝕊 *node of T, and let v*_1_ *and v*_2_ *be its children. Then we say that v is S*-consistent *if s*(*v*) ≠ *s*(*v*_1_) *and s*(*v*) ≠ *s*(*v*_2_).

*Furthermore, we say that T is S-consistent if each of its* 𝕊 *nodes is S-consistent*.

In the context of our work, the tree *T* is usually unknown, and only the orthology graph *G* is known. We use the existence/non-existence of a tree *T* corresponding to the relations in *G* to validate the orthology graph.

#### Definition 5

An orthology graph G is S-consistent if there exists a DS-tree T such that the two following conditions hold:

1. *ab* ∈ *E*(*G*) *if and only if a and b are orthologs in T;*
2. *T is S-consistent*.

The property of *S*-consistency has been investigated in depth in the recent years, and admits a convenient graph-theoretical characterization. First, we need to define a *triplet*. Let *T* be a tree, and let *x*,*y*,*z* ∈ *L*(*T*) be three distinct leaves. We say that *T* contains the triplet *xy*|*z* if *lca*_*T*_(*x*, *y*) < *lca*_*T*_(*x*,*z*) = *lca*_*T*_(*y*,*z*).

Theorem 6 (Lafond and El-Mabrouk (2014)). *An orthology graph G is S-consistent if and only if the two following conditions hold:*

1. *G is P*_4_-*free, i.e. has no path on 4 vertices without a chord;*
2. *for every x*,*y*,*z* ∈ *V*(*G*) *such that σ*(*x*), *σ*(*y*) *and σ*(*z*) *are distinct and such that xy*,*yz* ∈ *E*(*G*) but *xz* ∉ *E*(*G*), *we have that σ*(*x*)*σ*(*z*)|*σ*(*y*) *is a triplet of S*.

Therefore, *S*-consistent graphs can be characterized by the absence of *P*_4_’s and a set of forbidden *P*_3_’s given by the structure of the species tree. Given the set of primary orthology clusters, our goal is to infer the inter-cluster orthologs in a way that the resulting relations could be generated from a DAD-tree in a *S*-consistent manner.

### 2.3 The HyPPO orthology framework

We may now describe the main ingredients of our orthology inference framework. The flow of the pipeline is illustrated in Figure 2. Starting only from the DNA or amino acid sequences of a given set of genes (or proteins), our ultimate goal is to identify the cliques of primary orthologs, and the orthology relations between these cliques. Recall that this framework is modular, and that an implementation can be provided for each step individually. We first provide an outline of the tasks that each step must perform, and then proceed with the algorithmic details of how steps 2-4 were realized in the default implementation of our framework.

**Step 1.** We start by building a tentative *similarity graph*, in which two genes share an edge if they are “similar”. The pairwise similarity scores between the genes are typically used for this step. For instance, the graph may contain an edge *xy* iff the BLAST score between *x* and *y* is above a certain threshold. The edges can also be weighted by this score.

**Step 2.** As stated in Proposition 3, we expect the above similarity graph to contain only cliques. In practice, this might of course not always be the case and so in this step, we partition the graph into cliques of pairwise orthologous genes, which are assumed to be primary orthologs. Note that virtually every similarity-based inference method performs this task, although it should be observed that our requirement of pairwise orthology is strict, and no in-paralogs are allowed in the inferred groups.

**Step 3.** In order to obtain the inter-cluster relations in a *S*-consistent manner, a species tree *S* is needed. If known, this step can be omitted, but otherwise, a species tree can be inferred from the orthology clusters. Indeed, as we will show, some phylogenetic signal can be extracted from these clusters.

Note that in Figure 2, Step 3 points back to Step 2. This is because the framework allows reinferring the orthology clusters, but this time with the additional information of the species tree. The new clusters can then be used to reinfer the species tree, and this process can be looped until convergence. Although our framework supports such a loop, we did not use it in our experimentations, and leave this exploration for future work.

**Step 4.** In this final step, we assume that the orthology clusters and the species tree are known, but the orthologous gene pairs containing genes from distinct clusters are missing. Our goal is to recover these inter-cluster relations in a way to ensure *S*-consistency whilst preserving the properties of a DAD-tree. That is, we assume that in their history, two genes from two distinct clusters are separated by a divergent duplication edge, and that the predicted orthology relations can be explained by this type of history.

In the rest of this section, we describe how Steps 2, 3 and 4 were implemented in HyPPO.

#### 2.3.1 Prediction of orthology clusters

As partitioning a graph into cliques is a difficult algorithmic problem, most of the orthology prediction methods that perform clustering use a heuristic in order to deal with thousands, or even millions, of genes. In our case, we assume that genes are partitioned into families, which do not usually exceed a few hundred genes. In some cases, this makes it possible to use exact algorithms instead of heuristics. In fact, we provide two implementations for gene clustering: an exact method and a greedy heuristic. The exact method can be used on gene families with a reasonable number of genes (up to 200), and the heuristic for the other families.

The exact algorithm solves the cluster editing problem, which asks for the minimum number of edges to add or delete in a given graph so that every connected component of the resulting graph is a clique. We used the BBH graph, in which vertices are genes and edges are Bidirectional Best Hits (BBH), where two genes *x*, *y* form a BBH if *y* is the gene of *σ*(*y*) of maximum score with *x*, and vice-versa. The cluster editing problem is well-studied, and there are multiple methods that can find an optimal solution in an acceptable amount of time, including ILP formulations (Hartung and Hoos, 2015; Böcker *et al.*, 2011) and fixed-parameter algorithms (Böcker *et al.*, 2009). We used the ILP method (see (Böcker *et al.*, 2011) for details), ensuring that genes *a* and *b* from the same species would never belong to the same cluster by making the cost of adding the *ab* edge infinite.

The greedy heuristic proceeds in an agglomerative fashion using the pairwise scores. This can be seen as an adaptation of UPGMA with the restriction that clusters with a common species can never be joined. More precisely, at the start we treat each individual gene as a cluster. Then, we merge the two clusters *C*_1_ and *C*_2_ if *σ*(*C*_1_) ⋂ *σ*(*C*_2_) = ∅ and if they maximize max_a∈*C*_1_, *b*∈*C*_2__ *score*(*a*, *b*) (i.e. *C*_1_ and *C*_2_ have the two closest genes among all possible pairs of clusters). The algorithm stops when no more clusters can be merged. Note that we also tested other join criteria, for instance joining the two clusters that minimize the average distance as in UPGMA, but the above algorithm yielded the best results.

#### 2.3.2 Inference of a species tree from orthology clusters

We now show how orthology clusters may guide the construction of a species tree. In order to avoid confusion between species and clusters, we shall denote species with a *hat* symbol, e.g. *Ĉ* is a species. Consider the example provided in Figure 3. Here, the set of species *σ*(*C*_1_) appearing in cluster *C*_1_ forms a subset of the species *σ*(*C*_2_) from cluster *C*_2_. This suggests that *C*_1_ was given birth somewhere during the evolution of the *C*_2_ cluster. Moreover, the duplication that gave rise to *C*_1_ should have occurred in a species that existed at least as far back in time as the lca of *σ*(*C*_1_) in the true species tree *S*. This means that in the true gene tree for *C*_1_ ∪ *C*_2_, the duplication node *d* must satisfy *s*(*d*) ≥ *s*(*C*_1_) (where here *s* is with respect to the true species tree *S*). We argue that this suggests that the true species tree *S* contains the clade 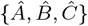, meaning that some node *u* of *S* has exactly 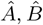 and 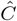 as its descendant leaves.

To justify this claim, observe that some species trees may require *losses* to explain the clusters, in a way that is analogous to gene tree and species tree *reconciliation* (see (Doyon *et al.*, 2011) for more details on this topic). Consider the two species tree *S*_1_ and *S*_2_ from Figure 3. If *S*_1_ is the true species tree and *d* is the duplication node that gave birth to the *C*_1_ cluster, then *s*(*d*, *S*_1_) ≥ *s*(*C*_1_, *S*_1_) = *x*, as argued in the previous paragraph. *T*_1_ is an example of an *S*_1_-consistent DAD-tree that satisfies this condition. If *S*_2_ is the true species tree, then *T*_1_ does not meet this requirement anymore. The tree *T*_2_ does satisfy *s*(*d*, *S*_2_) ≥ *s*(*C*_1_, *S*_2_) = *y*′. However, *s*(*C*_1_, *S*_2_) has 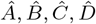 and 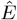 as its descendants, but 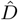 and 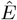 do not appear in *C*_1_. This can be explained by a gene loss prior to the ancestor of 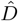 and 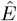, resulting in a loss in the 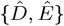 clade. The tree *T*_1_ does not require a loss to be explained, and therefore its corresponding species tree is preferred. Observe that any species tree that contains the 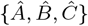 clade leads to a loss-less history, suggesting that this clade should be in the species tree.

Of course, the situation gets more difficult when *σ*(*C*_1_) is not a subset of *σ*(*C*_2_), or when there are more than two clusters. Below, we formalize the problem of reconstructing a species tree from for an arbitrary set of clusters while minimizing losses.

Let *C* be an orthology cluster. For convenience, we will make no distinction between *C* and its set of species *σ*(*C*), as this has no bearing on the inference procedure. For a species tree *S* and *x* ∈ *V*(*S*), the *clade* of *x* is defined as *clade*(*x*) = {*l* ∈ *L*(*S*): *l* is a descendant of *x*}. The set of clades of *S* is *clades*(*S*) = {*clade*(*x*): *x* ∈ *V*(*S*)}. For a given cluster *C*, let *S*_*C*_ be the subtree of *S* rooted at *lca*_*S*_(*C*). The number of losses of *C* with respect to *S*, denoted *l*_*S*_(*C*), is defined as the minimum number of clades to remove from *clades*(*S*_*C*_) so that the union of the remaining clades is exactly *C*. Another way to view *l*_*S*_(*C*) is as follows. Call a non-root node *v* of *S*_*C*_ *maximal C-free* if *clade*(*v*) ∩ *C* = ∅ but *clade*(*p*(*v*)) ∩ *C* ≠ ∅. Then *l*_*S*_(*C*) is the number of maximal *C*-free nodes in *S*_*C*_. As an example, the reader can verify in Figure 3 that *l*_*S*_2__ (*C*_1_) = 1 and *l*_*S*_2__ (*C*_2_) = 0. We may now formally define our species tree reconstruction problem.

The Species Tree Using Clusters (STUC) problem:

**Given**: a set of orthology clusters 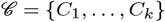.
**Find**: a species tree S that minimizes 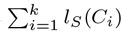.

Denote by 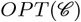 the minimum number of losses of a species tree for the STUC instance 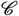. Note that 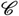 might not be sufficient to determine the complete species tree exactly, as in Figure 3 where any species tree containing the 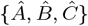 clade is optimal. However, the more clusters there are, the more precisely *S* can be inferred.

We do not know whether the STUC problem is NP-hard, although we suspect that is it. Nevertheless, we borrow ideas from (Mirarab *et al.*, 2014) in order to limit the space of possible species trees to those containing a predefined set of splits. For *X* ⊂ ∑, a *split* {*A*, *B*} for *X* is a partition of *X* into two non-empty subsets. We say that *S contains* the split {*A*, *B*} if there is a node *v* with children *v*_1_, *v*_2_ such that *A* = *clade*(*v*_1_) and *B* = *clade*(*v*_2_). Suppose that we are given a set of *allowable splits* 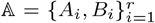, where for each *i*, *A*_*i*_ and *B*_*i*_ are disjoint subsets of ∑. Then we may ask for a species tree *S* that solves the STUC problem under the condition that every split contained in *S* appears in 𝔸. In the supplementary material, we show that this restricted version can be solved in polynomial time by dynamic programming, and then we discuss how a reasonable set 𝔸 can be constructed.

##### Theorem 7

*The STUC problem with species trees restricted to the allowable splits* 𝔸 *can be solved in time* 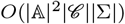.

#### 2.3.3 Inference of inter-cluster orthology relations

In this final step, we assume that the orthology clusters 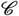 and the species tree *S* are known, but the orthologous gene pairs containing genes from distinct clusters are missing. Our goal is to recover these inter-cluster relations in a way to ensure *S*-consistency whilst preserving the properties of evolution under divergence after duplication. Note that unlike the last section, here we do not confound a cluster *C* and its set of species *σ*(*C*).

Consider the orthology clusters *C*_1_ and *C*_2_ in Figure 3. As we have argued in the previous section, the ancestral gene of *C*_1_ was born off a duplication node *d* satisfying *s*(*d*) ≥ *s*(*C*_1_). One implication of this is that if *σ*(*g*_2_) < *s*(*C*_1_), where *g*_2_ ∈ *C*_2_, then it is not be possible for a gene of *C*_1_ to be orthologous to *g*_2_. As an example, take the gene *a*_2_ ∈ *C*_2_ in Figure 3, and assume that *S*_1_ and *T*_1_ are the true species tree and gene tree, respectively. We have *σ*(*a*_2_) < *s*(*C*_1_),and so *a*_2_ is orthologous with no gene of *C*_1_. However, this is not true for *d*_2_ ∈ *C*_2_ for example, because *d*_2_ is a secondary ortholog with all of *C*_1_. This is because *σ*(*d*_2_) ⨁ *s*(*C*_1_).

In our implementation, when inferring inter-cluster relations between *C*_1_ and *C*_2_, we simply make as many pairs of genes orthologous whilst preserving *S*-consistency. As above, we also require that if *σ*(*g*_1_) < *s*(*C*_2_), then *g*_1_ is paralogous to all the genes in *C*_2_ (and vice-versa). This results in Algorithm 1. This works as described above in the case of *k* = 2 clusters. For larger *k*, we merge the clusters iteratively in a greedy manner.

More specifically, we first pick the cluster *C* for which *s*(*C*) is the the lowest in the species tree, breaking ties arbitrarily. We then ask: what other cluster *C*_*j*_ could have spawned the *C* cluster? We call *C*_*j*_ the *favorite cluster* of *C*, and assume that we are given a function 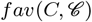 that finds the favorite cluster of *C* among a set of clusters 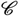. We then merge *C* with *C*_*j*_ by making all possible orthologies - namely, we make *c*_*j*_ ∈ *C*_*j*_ orthologous to every gene of *C* iff *σ*(*c*_*j*_) ⨁ *s*(*C*). Afterwards, *C*_*j*_ is redefined as *C*_*j*_ ∪ *C*, *C* is emptied and we proceed with the next cluster. We show that Algorithm 1 preserves *S*-consistency as desired.

##### Algorithm 1 Algorithm to infer inter-cluster orthologs

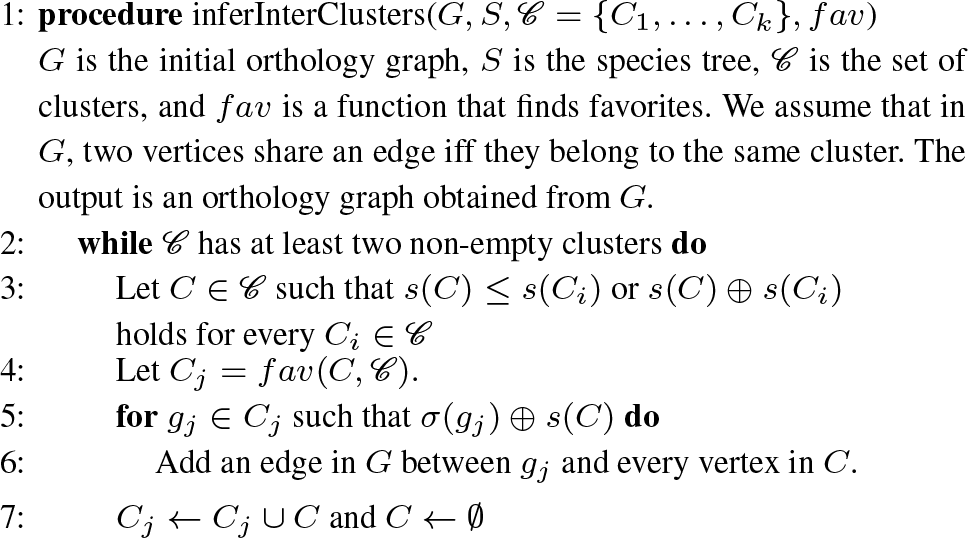

##### Theorem 8

The orthology graph obtained from Algorithm 1 is S- consistent.

In our implementation, we used the following simple *fav* function: for each *x* ∈ *C*, find the gene *y* ∈ Γ \ *C* of maximum score. Then *x* votes for the cluster that contains *y*. The favorite cluster of *C* is then the one with the highest number of votes. Only the members of the genes initially in *C*, as given in the input, are allowed to vote.

## 3 Results

We compare the default implementation of the HyPPO pipeline with two other orthology prediction methods: OMA-GETHOGS and OrthoMCL. To our knowledge, OMA-GETHOGS is the method that is the most similar to ours, as it outputs both the 1-to-1 orthology groups, and the pairwise orthology relations. The groups can be interpreted as the primary orthologs. As for OrthoMCL, it is a similarity-based clustering method that focuses on grouping orthologs, and so it is not expected to find all the secondary orthologs. Nevertheless, we include it for the sake of comparison, as OrthoMCL is one of the most popular orthology prediction tools. HyPPO was tested under two settings: one in which the species tree is unknown, and the other in which the true simulated species tree is provided. OMA-GETHOGS was provided the true species tree in all experiments [^1^ The program does not require a species tree as input, but OMA-GETHOGS produced an error message when not given one - for reasons that are yet to be determined. Nevertheless, better results are expected when the true species tree is known.].

We performed our experiments on both simulated and real empirical datasets. In the simulated datasets, the true orthologs and the primary orthologs are known. This is not true for the real datasets. As in (Altenhoff *et al.*, 2013), we used a set of gold standard, manually curated, gene trees from the SwissTree database (Wapinski *et al.*, 2007). These trees are annotated with duplication and speciation events, and so the underlying orthology/paralogy relations can be considered as “gold standard” relations. We did not, however, include the analysis of the primary vs secondary orthologs, as those are not determined in the real datasets. Note that in both simulated and real datasets, the genes were already separated into homologous families. All the programs were executed on the separated gene families, thereby avoiding predictions between non-homologous genes. For HyPPO, the sequences were aligned with MAFFT (Katoh *et al.*, 2002), and the similarity values for the first step were obtained from the pairwise identity percentages computed from these alignments (number of identical base pairs divided by the length of the longest sequence, not counting gaps).

### 3.1 Performance metrics

In our experiments, we address two questions: is the set of all pairwise relations predicted correct? And, can the method distinguish the primary orthologs from the secondary orthologs? For the pairwise relations, we used precision, recall and accuracy as our performance metrics. The precision *Prec* and recall *Rec* values are calculated from the orthology relations as follows: 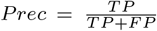 and 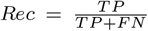. Here, *TP* means True Positive (the number of relations correctly predicted as orthology), *FP* means False Positive (a paralogy relation incorrectly predicted as orthology) and *FN* means False Negative (an orthology relation left undetermined or predicted as paralogy). As for accuracy, it consists in the total number of relations predicted correctly (including both orthologs and paralogs) divided by 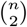, the total number of relations [^2^ Note that Ortho-MCL does not predicts paralogs, aside from the recent ones. In our computation of accuracy, we interpreted as paralogy the set relations not predicted as orthology - otherwise, leaving these relations as undetermined made the accuracy too low for comparison.].

In the simulated datasets containing divergent duplication edges, we also evaluated the ability to recover the primary orthologs. The metric we used is the *cluster score* defined as follows: let *P*_1_ and *P*_2_ be two partitions of a given set of elements. Here, *P*_1_ would be the predicted primary orthologs, and *P*_2_ the “true” simulated primary orthologs. Take a graph in which the vertices are *P*_1_ ∪ *P*_2_ (i.e. one vertex for each subset), add an edge *X*_1_ *X*_2_ between each pair *X*_1_ ∈ *P*_1_, *X*_2_ ∈ *P*_2_ and weigh this edge by |*X*_1_ ∩ *X*_2_|. Then the cluster score between *P*_1_ and *P*_2_ is the value of a maximum matching in this graph, divided by the number of genes. Hence the cluster score is 1 iff *P*_1_ and *P*_2_ are equal. The idea behind this score is that this value yields the proportion of elements from *P*_1_ that must be moved from one set to another in order to obtain *P*_2_ (if the score is *c*, then a fraction of 1 − *c* elements from *P*_1_ need to be moved).

### 3.2 Simulated datasets

For our simulations, we used SimPhy (Mallo *et al.*, 2015) to generate a set of species trees along with a set of gene trees evolving within these species trees. As SimPhy incorporates speciation and duplication events in its simulations, the true sets of orthologs and paralogs are known. We generated 40 species trees and 400 gene trees - 10 gene families for each species tree. The number of species in each species tree was chosen uniformly at random between 30 and 50. The gene trees were subject to the events of speciation, duplication and losses. We then used the INDELiBLE (Fletcher and Yang, 2009) module provided with SimPhy to simulate the evolution of gene sequences on each gene tree. The nucleotide evolutionary model was selected at random for each gene family, each evolving under a General-Time Reversible model (GTR) with rates sampled from a 6-dimensional Dirichlet distribution. The only input to the programs was the set of extant gene sequences and, depending on the method, the true simulated species tree.

In order to evaluate the impact of the rates of duplications, losses and substitutions, we used four sets of parameters for the simulations, yielding four datasets named as follows: Standard, Eventful, Fast and Slow. Standard uses default parameters, Eventful has an increased rate of duplications and losses, Fast has high mutation rates and Slow has low mutation rates. For each dataset, 10 species trees were generated. In both the Standard and Eventful datasets, the tree-wide substitution rate was set to 0.000005 (the default value in SimPhy). In the Default dataset, the duplication and loss rates were set to 0.0000005 each, and to 0.000001 in the Eventful dataset. Higher duplication rates led to trees with too many genes in the trees (averaging in the thousands, making our analysis too computationally intensive) while lower values yielded gene trees with too few paralogs. In the Fast dataset, the duplication/loss rates were set as in the Standard setting, but the tree-wide substitution rate was increased to 0.00005, In the Slow dataset, this parameter took value 0.0000005. The Fast dataset led to difficulties in alignments, due to the presence of many substitutions and very large gaps, whereas the Slow dataset had a small amount of substitutions, resulting in sequences with fewer differences.

These simulations do not consider the phenomenon of divergence after duplication. That is, the children of 𝔻 nodes in the generated gene trees do not differentiate from each other, and both copies continue to evolve under the same model. Therefore we also simulated divergent edges by choosing, for every 𝔻 node in the simulated gene trees, one child edge arbitrarily and multiplying its length by some factor *ℓ*. Thus for each gene tree and each *ℓ* ∈ {1, 2, 8, 50}, we used INDELiBLE to generate a set of extant gene sequences obtained by multiplying the length each divergent edge by *ℓ*. Each gene family gave rise to four sets of sequences, resulting in a total of 1600 gene families to analyze. In the cases where *ℓ* > 1, the primary orthologs can be deduced from the divergent duplication edges.

The average accuracy, precision, recall and cluster scores, taken over the 1600 simulated gene families, are presented in Table 1. Figure 4 shows the average accuracy per gene family, for each dataset and possible value of *ℓ*. HyPPO outperforms the other methods on every dataset, even when not given a species tree. Moreover, the accuracy of HyPPO always improves when the true species tree is known. We observe that the accuracy of OMA tends to decrease as *ℓ* is increased, whereas for HyPPO and OrthoMCL, the impact of changing *ℓ* seems to vary.

**Fig. 4:**
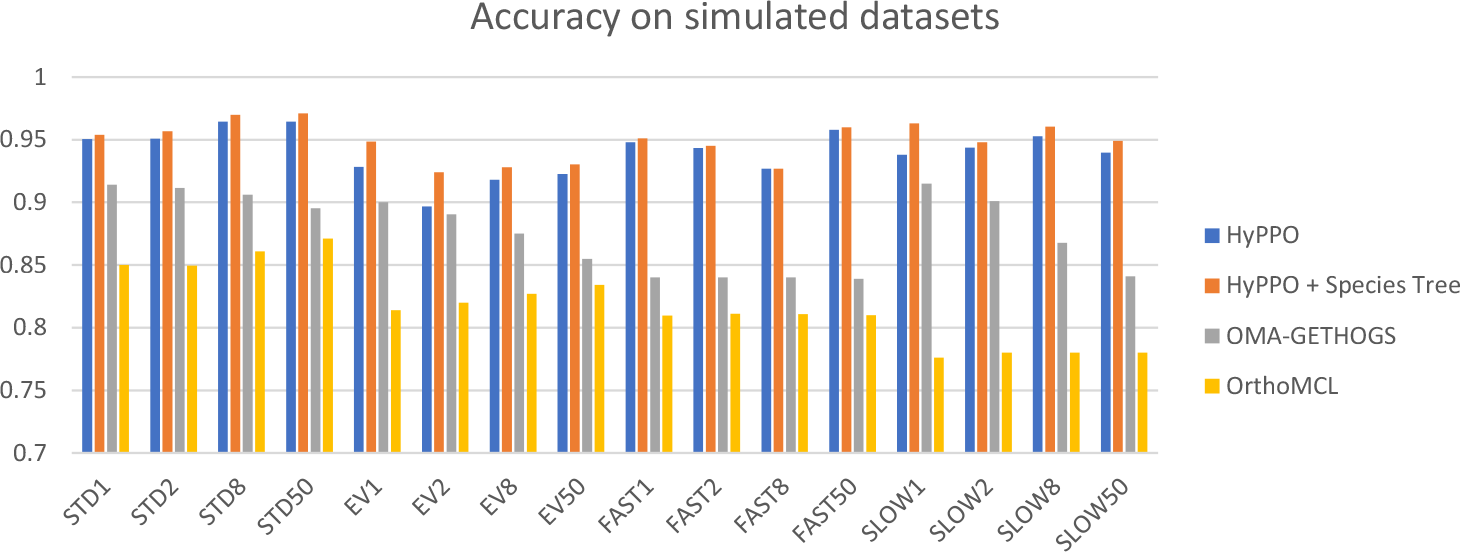
Accuracy of the methods for each dataset and each branch multiplication factor *ℓ* ∈ {1,2,8,50}. STD stands for Standard and EV for Eventful.

Figure 5 shows the average cluster scores per dataset. Note that the scores for the trees having multiplication factor *ℓ* = 1 are not presented, as in these cases the primary orthologs cannot be determined with certainty. Here again, we see that HyPPO is better at identifying the primary orthologs. Observe that knowledge of the species tree is inconsequential here, as the species tree does not impact the gene clustering step in our pipeline. We also observe that for all methods, the best cluster scores are achieved at *ℓ* = 50, which is expected since this creates a greater separation between the primary and secondary orthologs.

**Fig. 5:**
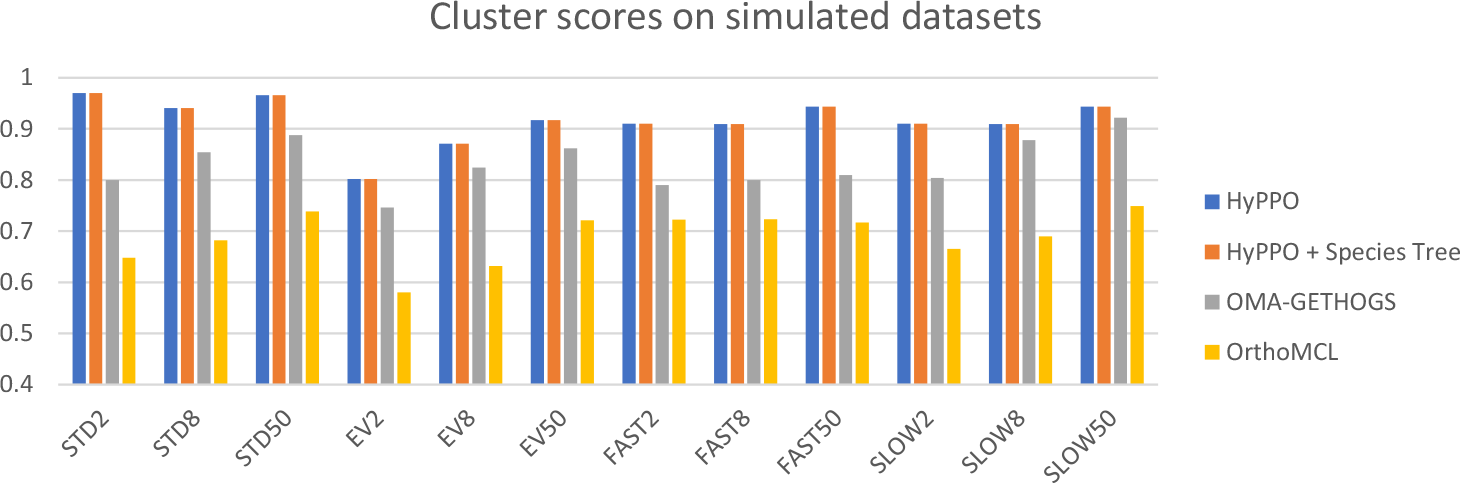
Cluster scores of the methods for each dataset and each branch multiplication factor *ℓ* ∈ {1,2,8,50}.

The comparison of precision and recall is shown in Figure 7 for *ℓ* = 8. The other values of *ℓ* yielded similar results. One can see that OMA typically attains better precision values. This is at the cost of a much lower recall, as HyPPO outperforms the other methods in this aspect. Therefore, the orthologs predicted by OMA are truly orthologs slightly more often, but the method tends to miss more of the orthology relations.

### 3.3 Empirical datasets

We used 8 gene trees from the gold standard SwissTree database. This included the Popeye domain family (POP), the NOX ‘ancestral-type’ subfamily NADPH oxidases (NOX), the V-type ATPase beta subunit (VATB), the Serine incorporator family (SERC), the Sulfatase-modifying factor 1 family (SUMF), the HOX cluster genes family 9-14 (HOX), the Asterix family (ARX) and the Cited family (CITE). These gene families were chosen because they belong to Eukaryotes and, according to the curators, are not suspected to have undergone horizontal gene transfer.

Figure 6 shows the accuracy of each method for each gene tree. Contrary to the simulated datasets, HyPPO does not outperform OMA on every single dataset, as it performs poorly on the VATB and SERC families. Notably, Ortho-MCL attains almost perfect accuracy on the VATB family (0.96) whereas HyPPO is not much more than half as accurate (0.59). The reason appears to be that most gene pairs are orthologous (95.7%), since the duplications are low in this tree. HyPPO infers multiple orthology clusters joined together by duplications, resulting in many false paralogs (we predict 47% pairs as orthologous), whereas OrthoMCL puts almost all the genes into one big orthology cluster, making its predictions more accurate. A similar phenomenon appears to occur with SERC. Nevertheless, HyPPO achieves similar or superior accuracy on the other trees, and so it remains competitive. Observe that as in the simulations, when HyPPO is given the species tree (provided by SwissTree), the accuracy increases.

**Fig. 6:**
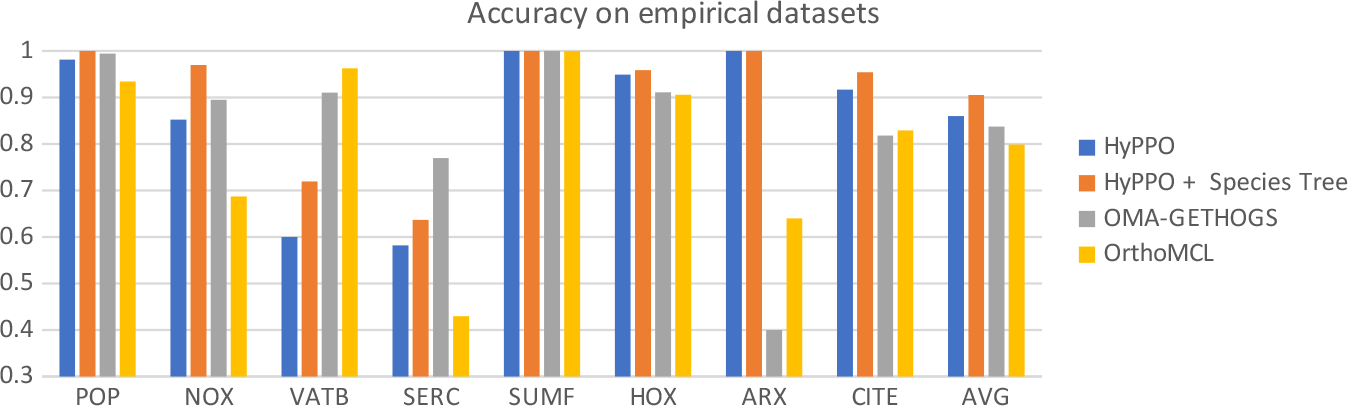
Accuracy of the methods for each tree in our empirical dataset. The last column shows the average accuracy of the methods.

**Fig. 7:**
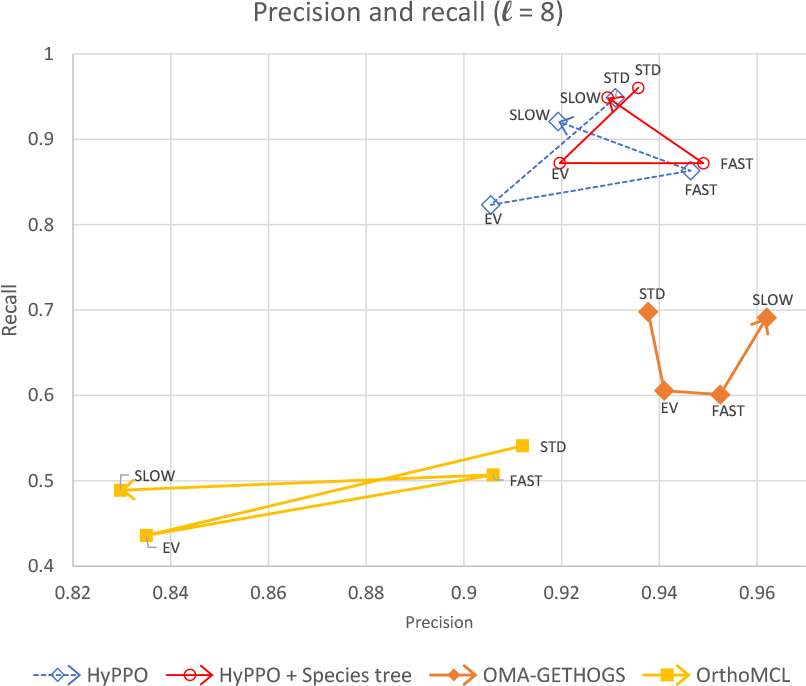
Precision versus recall values of each method and each dataset for *ℓ* = 8. Each method has four corresponding points, one for each dataset, and are ordered as follows: Standard - Eventful - Fast - Slow. For example the second point of each path for the analysis of the Eventful dataset.

As for precision and recall, the four methods attained similar values. Once again, OMA offers slightly better precision at the cost of lower recall. The averages over all 8 trees are presented in Table 2.

**Table 2.** Average accuracy, precision and recall over the set of 8 trees from SwissTree.

|  | Accuracy | Precision | Recall |
| --- | --- | --- | --- |
| HyPPO | .860 | .941 | .785 |
| HyPPO + Species Tree | .905 | .945 | .881 |
| OMA-GETHOGS | .837 | .950 | .711 |
| OrthoMCL | .800 | .859 | .680 |

## 4 Discussion

Although the results of HyPPO are promising, there is still room for improvements. Each step of the pipeline could be implemented in a more sophisticated manner, and the DAD model around which HyPPO is built could be more realistic. When a duplication node is not followed by accelerated mutations, there is no special post-duplication divergent edge, and it could be argued that the orthology clusters should contain in-paralogs. In the future, we might thus consider merging the clusters that are share a high enough degree of similarity. We have also ignored the effects of convergent evolution, even though it is possible that two independent divergent duplication events lead to similar genes. If this occurs, one cannot guarantee that similar genes are orthologs, and that primary orthologs form cliques.

On the practical side, we assume that the genes are already partitioned into homologous families, which allows the computation of a multiple sequence alignement and eases the clustering phase. Future versions of the framework should be able to predict orthologs and paralogs from whole proteomes, as OrthoMCL and OMA-GETHOGS do. However, one could also argue that partitioning genes into homologous families and predicting orthology are two different tasks. The aforementioned programs perform both simultaneously. A better approach might be to use methods dedicated to homology clustering, and then refine the orthology/paralogy relations within each family using HyPPO.

Also, the reconstruction of a species tree from the clusters is the main bottleneck of HyPPO in terms of time. To give a rough idea, in the worst case, some datasets resulted in a set of about 3000 allowable splits, which took our algorithm 4-5 hours to solve. This motivates the need to improve the complexity of the dynamic programming formulation given in this paper. Also, in the near future we plan to evaluate the quality of the predicted species tree. Note however that our current algorithm may take many arbitrary decisions, and we must refine this process before performing a careful species tree analysis. Finally, it will be interesting to include a wider range of orthology prediction software in a future comparative study, and to establish a procedure to analyze primary orthologs on real datasets - for instance by predicting these orthologs by comparing the ratio of branch lengths following a duplication.

## 5 Conclusion

In this work, we have introduced the notion of orthology in the presence of divergence after duplication and have addressed the problem that some orthologs may behave as paralogs from a functional point of view. We therefore propose to make a distinction between the orthologs that have or have not been separated by a divergent duplication event. As we have shown, the HyPPO framework is able to distinguish paralogs, primary orthologs and secondary orthologs with better accuracy than its competitors. This is true even if divergence after duplication does not always occur. Our work is far from over though, as HyPPO offers many opportunities to improve these results even further. Notably, it remains to evaluate alternative algorithms for each step in our pipeline, and to identify which ones yield the best results. Finally, our work raises some interesting questions from a theoretical standpoint. The complexity of inferring a species tree from gene clusters is left open, and it remains to evaluate how many clusters are necessary to reconstruct a high quality species tree.

## References

Altenhoff, A. M. and Dessimoz, C. (2012). Inferring orthology and paralogy. Evolutionary Genomics: Statistical and Computational Methods, pages 259–279.

Altenhoff, A. M., Studer, R. A., Robinson-Rechavi, M., and Dessimoz, C. (2012). Resolving the ortholog conjecture: orthologs tend to be weakly, but significantly, more similar in function than paralogs. PLoS Computational Biology, 8(5).

Altenhoff, A. M., Gil, M., Gonnet, G. H., and Dessimoz, C. (2013). Inferring hierarchical orthologous groups from orthologous gene pairs. PLoS ONE, 8(1).

Böcker, S., Briesemeister, S., Bui, Q. B. A., and Truß, A. (2009). Going weighted: Parameterized algorithms for cluster editing. Theoretical Computer Science.

Böcker, S., Briesemeister, S., and Klau, G. W. (2011). Exact algorithms for cluster editing: Evaluation and experiments. Algorithmica, 60(2), 316–334.

Cardoso-Moreira, M., Arguello, J. R., Gottipati, S., Harshman, L. G., Grenier, J. K., and Clark, A. G. (2016). Evidenceforthe fixationofgeneduplicationsbypositive selection in drosophila. Genome Research, 26(6), 787–798.

Chen, X. and Zhang, J. (2012). The ortholog conjecture is untestable by the current gene ontology but is supported by rna sequencing data. PLoS Computational Biology, 8(11), e1002784.

Dondi, R., Lafond, M., and El-Mabrouk, N. (2017a). Approximating the correction of weighted and unweighted orthology and paralogy relations. Algorithms for Molecular Biology, 12(1), 4.

Dondi, R., Mauri, G., and Zoppis, I. (2017b). Orthology correction for gene tree reconstruction: Theoretical and experimental results. Procedia Computer Science.

Doyle, M. A., Gasser, R. B., Woodcroft, B. J., Hall, R. S., and Ralph, S. A. (2010). Drug target prediction and prioritization: using orthology to predict essentiality in parasite genomes. BMC Genomics, 11(1), 222.

Doyon, J.-P., Ranwez, V., Daubin, V., and Berry, V. (2011). Models, algorithms and programs for phylogeny reconciliation. Briefings in Bioinformatics, 12(5).

Emms, D. M. and Kelly, S. (2015). Orthofinder: solving fundamental biases in whole genome comparisons dramatically improves orthogroup inference accuracy. Genome Biology, 16(1), 157.

Fitch, W. M. (2000). Homology: a personal view on some of the problems. Trends in Genetics, 16(5), 227–231.

Fletcher, W. and Yang, Z. (2009). Indelible: a flexible simulator of biological sequence evolution. Molecular Biology and Evolution, 26(8), 1879–1888.

Gabaldón, T. and Koonin, E. V. (2013). Functional and evolutionary implications of gene orthology. Nature Reviews Genetics, 14(5), 360–366.

Hartung, S. and Hoos, H. H. (2015). Programming by optimisation meets parameterised algorithmics: A case study for cluster editing. In Int. Conference on Learning and Intelligent Optimization, pages 43–58. Springer.

Hellmuth, M., Hernandez-Rosales, M., Huber, K. T., Moulton, V., Stadler, P. F., and Wieseke, N. (2013). Orthology relations, symbolic ultrametrics, and cographs. Journal of Mathematical Biology, pages 1–22.

Hellmuth, M., Wieseke, N., Lechner, M., Lenhof, H.-P., Middendorf, M., and Stadler, P. F. (2015). Phylogenomics with paralogs. Proceedings of the National Academy of Sciences, 112(7), 2058–2063.

Hernandez-Rosales, M., Hellmuth, M., Wieseke, N., Huber, K. T., Moulton, V., and Stadler, P. F. (2012). From event-labeled gene trees to species trees. BMC Bioinformatics, 13(19), S6.

Innan, H. and Kondrashov, F. (2010). The evolutionofgene duplications: classifying and distinguishing between models. Nature Reviews Genetics, 11(2), 97.

Jones, M., Paul, C., and Scornavacca, C. (2016). On the consistency of orthology relationships. BMC Bioinformatics, 17(14),416.

Jordan, I. K., Wolf, Y. I., and Koonin, E. V. (2004). Duplicated genes evolve slower than singletons despite the initial rate increase. BMC Evolutionary Biology, 4(1).

Jothi, R., Zotenko, E., Tasneem, A., and Przytycka, T. M. (2006). Coco-cl: hierarchical clustering of homology relations based on evolutionary correlations. Bioinformatics, 22(7), 779–788.

Katoh, K., Misawa, K., Kuma, K.-i., and Miyata, T. (2002). Mafft: a novel method for rapid multiple sequence alignment based on fast fourier transform. Nucleic Acids Research, 30(14), 3059–3066.

Koonin, E. V. (2005). Orthologs, paralogs, and evolutionary genomics. Annual Review of Genetics, 39, 309–338.

Kristensen, D. M., Wolf, Y. I., Mushegian, A. R., and Koonin, E. V. (2011). Computational methods for gene orthology inference. Briefings in Bioinformatics.

Lafond, M. and El-Mabrouk, N. (2014). Orthology and paralogy constraints: satisfiability and consistency. BMC Genomics, 15(6), S12.

Lafond, M., Dondi, R., and El-Mabrouk, N. (2016). The link between orthology relations and gene trees: a correction perspective. Algorithms for Molecular Biology, 11(1), 4.

Lechner, M., Findeiß, S., Steiner, L., Marz, M., Stadler, P. F., and Prohaska, S. J. (2011). Proteinortho: detection of (co-) orthologs in large-scale analysis. BMC Bioinformatics, 12(1), 124.

Li, L., Stoeckert, C. J., and Roos, D. S. (2003). Orthomcl: identification of ortholog groups for eukaryotic genomes. Genome Research, 13(9), 2178–2189.

Lynch, M. and Conery, J. S. (2000). The evolutionary fate and consequences of duplicate genes. Science, 290(5494), 1151–1155.

Mallo, D., de Oliveira Martins, L., and Posada, D. (2015). Simphy: Phylogenomic simulation of gene, locus, and species trees. Systematic Biology, 65(2), 334–344.

Mirarab, S., Reaz, R., Bayzid, M. S., Zimmermann, T., Swenson, M. S., and Warnow, T. (2014). Astral: genome-scale coalescent-basedspeciestree estimation. Bioinformatics, 30(17), i541–i548.

Nehrt, N. L., Clark, W. T., Radivojac, P., and Hahn, M. W. (2011). Testing the ortholog conjecture with comparative functional genomic data from mammals. PLoS Computational Biology, 7(6), e1002073.

O’brien, K. P., Remm, M., and Sonnhammer, E. L. (2005). Inparanoid: a comprehensive database of eukaryotic orthologs. Nucleic Acids Research, 33.

Powell, S., Szklarczyk, D., Trachana, K., Roth, A., Kuhn, M., Muller, J., Arnold, R., Rattei, T., Letunic, I., Doerks, T., et al. (2011). eggnog v3. 0: orthologous groups covering 1133 organisms at 41 different taxonomic ranges. Nucleic Acids Research, 40(D1).

Roth, A. C., Gonnet, G. H., and Dessimoz, C. (2008). Algorithm of oma for large-scale orthology inference. BMC Bioinformatics, 9(1), 518.

Soria, P. S., McGary, K. L., and Rokas, A. (2014). Functional divergence for every paralog. Molecular Biology and Evolution, 31(4), 984–992.

Stolzer, M., Lai, H., Xu, M., Sathaye, D., Vernot, B., and Durand, D. (2012).Inferring duplications, losses, transfers and incomplete lineage sorting with nonbinary species trees. Bioinformatics, 28(18), i409–i415.

Studer, R. A. and Robinson-Rechavi, M. (2009). How confident can we be that orthologs are similar, but paralogs differ? Trends in Genetics, 25(5), 210–216.

Swenson, K. M. and El-Mabrouk, N. (2012). Gene trees and species trees: irreconcilable differences. BMC Bioinformatics, 13(19), S15.

Tatusov, R. L., Fedorova, N. D., Jackson, J. D., Jacobs, A. R., Kiryutin, B., Koonin, E. V., Krylov, D. M., Mazumder, R., Mekhedov, S. L., Nikolskaya, A. N., et al. (2003). The cog database: an updated version includes eukaryotes. BMC Bioinformatics, 4(1), 41.

Thomas, P. D., Wood, V., Mungall, C. J., Lewis, S. E., Blake, J. A., Consortium, G. O., et al. (2012). On the use of gene ontology annotations to assess functional similarity among orthologs and paralogs: a short report. PLoS Computational Biology, 8(2), e1002386.

Train, C.-M., Glover, N. M., Gonnet, G. H., Altenhoff, A. M., and Dessimoz, C. (2017). Orthologous matrix (oma) algorithm 2.0: more robust to asymmetric evolutionary rates and more scalable hierarchical orthologous group inference. Bioinformatics, 33(14), i75–i82.

Ullah, I., Sjöstrand, J., Andersson, P., Sennblad, B., and Lagergren, J. (2015). Integrating sequence evolution into probabilistic orthology analysis. Systematic Biology, 64(6), 969–982.

Van der Heijden, R. T., Snel, B., Van Noort, V., and Huynen, M. A. (2007). Orthology prediction at scalable resolution by phylogenetic tree analysis. BMC Bioinformatics, 8(1), 83.

Wapinski, I., Pfeffer, A., Friedman, N., and Regev, A. (2007). Automatic genome-wide reconstructionofphylogeneticgene trees. Bioinformatics, 23(13), i549–i558.

Woods, S., Coghlan, A., Rivers, D., Warnecke, T., Jeffries, S. J., Kwon, T., Rogers, A., Hurst, L. D., and Ahringer, J. (2013). Duplication and retention biases of essential and non-essential genes revealed by systematic knockdown analyses. PLoS Genetics, 9(5), e1003330.

Zhang, J. (2003). Evolution by gene duplication: an update. Trends in Ecology & Evolution, 18(6), 292–298.

Zheng, C., Swenson, K., Lyons, E., and Sankoff, D. (2011). Omg! orthologs in multiple genomes-competing graph-theoretical formulations. In WABI.

